# Undergraduate research project uncovers a novel strain of amphibian chytrid fungus in the Northeastern United States

**DOI:** 10.1101/2024.11.21.624705

**Authors:** Katharine York, Mariah Mitchell, Tyler Cardoso, Ashley Chapdelaine, Yuhui Song, M. Catherine Duryea

## Abstract

Investigation of the genetic variation of *Batrachochytrium dendrobatidis* (*Bd* or chytrid fungus) in New Hampshire (NH) in the northeastern U.S. has shown the presence of genetic mutations which are currently undocumented in the literature. DNA was collected as part of a long-term monitoring project in New Hampshire to test for the presence of *Bd* in amphibian populations. In this monitoring project, we are detecting the presence of *Bd* in multiple populations, but the observed amphibians do not appear to have increased mortality or other symptoms related to the fungus. To further investigate the strain of *Bd* affecting NH amphibians, we sequenced samples that were collected in 2019, as part of your yearly monitoring effort. Utilizing the Basic Local Alignment Search Tool (BLAST) and sequences available on GenBank, we found that there are two different strains of chytrid in the observed NH amphibians. These strains are similar to strains noted previously in New England, but exhibit unique point mutations and in one case, a deletion. These variations could affect the impact that *Bd* has on amphibians in this region, and therefore could contribute to a better understanding of the variation in strains of chytrid and how they impact amphibian populations around the world. This study highlights the importance of monitoring efforts conducted at smaller, non-research institutions and we encourage other small institutions to take up similar monitoring efforts for *Bd* or other national or global conservation concerns.

## Introduction

Amphibian populations are declining around the world due to *Batrachochytrium dendrobatidis* (henceforth *Bd* or chytrid fungus) and the disease it causes, Chytridiomycosis (Lips *et al*., 2006; Skerratt *et al*., 2007). *Bd* is a zoosporic fungus that affects the keratinized epidermal cells of amphibians through irregular epidermal hyperplasia and hyperkeratosis (Morehouse *et al*., 2002). Amphibians are known bioindicators, meaning they are important early warning signs of issues and concerns in ecosystems. There are multiple hypotheses accepted among researchers as to what where *Bd* originated, how *Bd* was introduced to frogs globally, and why it is causing declines in many amphibian populations (Rosenblum *et al*., 2010; Farrer *et al*., 2011; Fisher & Garner, 2020). However, as *Bd* becomes increasingly widespread, we are still attempting to describe the geographic extent and genetic diversity of this fungus.

Previous studies have indicated that the genetic strain of *Bd* can play a role in how it will affect amphibians. In a study by Gahl *et al*., 2012, several species of amphibians were exposed to two different strains of *Bd*. The two strains were noted as a northeastern strain (JEL404) and a strain that caused die-offs of amphibians in Panama (JEL423) (Gahl *et al*., 2012). Bullfrogs (*Lithobates catesbeianus*) and spring peepers (*Pseudacris crucifer*) were observed to be unaffected by *Bd* exposure from either strain. However, wood frogs (*Lithobates sylvaticus*) exhibited mortality from both *Bd* strains, and green frogs (*Lithobates clamitans*) had complete mortality from JEL423 but no mortality from JEL404 (Gahl *et al*., 2012).

Genetic variation in different strains of *Bd* is known to be a large influence on the variation in virulence among strains of *Bd*. Through testing of passages (hereditary passing of genetic information over generations), it was shown that the more passages there are for a strain of *Bd*, the less virulent it becomes (Refsnider *et al*., 2015). In Refsnider *et al*., (2015), they compared JEL427-P9 zoospores that had 9 passages to JEL427-P39 zoospores that had 39 passages. JEL427-P9 was shown to have an increased mortality of frogs when compared with JEL427-P39. Zoospores were also created at a greater rate for the strain with only 9 passages versus the 39-passage strain (Refsnider *et al*., 2015). Recent work has also shown that gene expression patterns vary greatly both within and between lineages of *Bd* (McDonald *et al*., 2020). These studies indicate that chytrid strain variation can affect the virulence of *Bd* and highlight the importance of documenting chytrid fungal strain variation to better understand how it affects amphibian populations.

In this article, we report on the findings of an ongoing undergraduate research project to monitor frogs for chytrid fungus in central and southern New Hampshire (NH). Amphibian chytrid fungus has been detected in nearby regions of the northeastern U.S., including Maine and New York (Longcore *et al*., 2007; Lenker *et al*., 2014). We also examine the genetics of *Bd* to see if there is variation in the strain affecting the central and southern New Hampshire area. Discovering genetic variation could provide evidence that genetics are playing a part in virulence and mortality rates of the amphibian chytrid fungus affecting NH populations, and therefore call for further investigation of these fungal strains. In this article, we address this question, *e*.*g*.,: *Is the NH strain of Bd different from strains found in other parts of the world?* There are three hypotheses for this study experiment: First, there may be genetic changes that cause the NH strain to be locally adapted, or affect the virulence of the fungus. Second, there could be variable sites that are silent mutations and have no effect on the fungus or its virulence. Lastly, there could be little to no variable sites, and the *Bd* strain found in NH populations is identical to that found in other populations.

Additionally, we report on the results for a *Bd* monitoring effort conducted in 2019. Through this effort, we sampled several ponds in the central and southern New Hampshire area. Our monitoring involves undergraduate research training and outreach to community members. Here, we report on the percentage of frogs that tested positive at these locations and give a template for how other small institutions can help in the *Bd* monitoring effort.

## Materials and methods

### Field Collections

Amphibians were examined at eight collection sites across central and southern New Hampshire (Table 1, Fig. 1). Samples were collected from late July through October of 2019 (Appendix 1). Our efforts to obtain samples for *Bd* analysis focused on being able to collect samples accurately as well as humanely for the amphibians. Frogs were captured at local water bodies using small-holed nets and then temporarily housed in 5-gallon buckets with secured lids. Buckets were filled approximately halfway with water from the same source amphibians were being collected from, and nearby sticks were placed in the buckets to provide a space for the frogs to perch above water. Once enough specimens were captured, field efforts moved into field data collection and tissue swabbing. At that time no additional specimens were caught to eliminate the possibility of duplicate sampling. Each frog was handled with a new pair of disposable nitrile gloves to reduce cross contamination and assigned an individual sample identification number (Fig. 2). Data was recorded for each amphibian including: date of collection, time of collection, species, sex (when it could be determined, based on external features), age, waterbody name, type of water (natural pond, drainage pond, wetland, etc.), waterbody town location, water temperature, visible signs of chytrid, and any other relevant observations to amphibian health (Appendix 1). Visual documentation photographs were taken of each specimen’s dorsal and ventral side with the specimen ID number included for reference (Fig. 2).

**Table 1:** Results of a *Bd* monitoring effort conducted in 2019 in several towns across New Hampshire. Species abbreviations: BUF = Bullfrog, GRF = Green Frog, SPP = Spring Peeper, WOF = Wood Frog, PIF = Pickerel Frogs. For each species, the first column (Tot.) represents the total frogs collected and the second column (Pos.) represents the number of frogs that were positively identified to have *Bd*. The last two columns represent the total number of frogs that tested positive (Pos.) and negative (Neg.) for *Bd* at each collection site.

| Town | Site | Lat. | Long. | BUF | BUF | GRF | GRF | SPP | SPP | WOF | WOF | PIF | PIF | Sum | Sum |
| --- | --- | --- | --- | --- | --- | --- | --- | --- | --- | --- | --- | --- | --- | --- | --- |
|  |  |  |  | Tot. | Pos. | Tot. | Pos. | Tot. | Pos. | Tot. | Pos. | Tot. | Pos. | Pos. | Neg. |
| Chichester | Lynxfield Pond | 43.27 | -71.41 |  |  | 3 | 1 |  |  |  |  |  |  | 1 | 2 |
| Chichester | Marsh Pond | 43.24 | -71.39 |  |  | 3 | 3 |  |  |  |  | 4 | 3 | 6 | 1 |
| Concord | Carter Hill Orchard | 43.23 | -71.61 | 4 | 4 | 2 | 1 |  |  |  |  |  |  | 5 | 1 |
| Manchester | Dorrs Pond | 43.01 | -71.45 | 4 | 3 | 16 | 11 |  |  |  |  |  |  | 14 | 6 |
| Pembroke | Hill School | 43.16 | -71.47 |  |  | 5 | 3 |  |  |  |  |  |  | 3 | 2 |
| Pembroke | House Pond | 43.16 | -71.48 | 4 | 4 | 8 | 6 |  |  |  |  |  |  | 10 | 2 |
| Pembroke | Melissa Drive | 43.16 | -71.48 |  |  | 1 | 0 | 2 | 0 | 4 | 2 |  |  | 2 | 5 |
| Pembroke | Pembroke Academy | 43.15 | -71.45 | 9 | 6 | 5 | 1 |  |  |  |  |  |  | 7 | 7 |
| **Total** |  |  |  |  |  |  |  |  |  |  |  |  |  | **48** | **26** |

**Figure 1:**
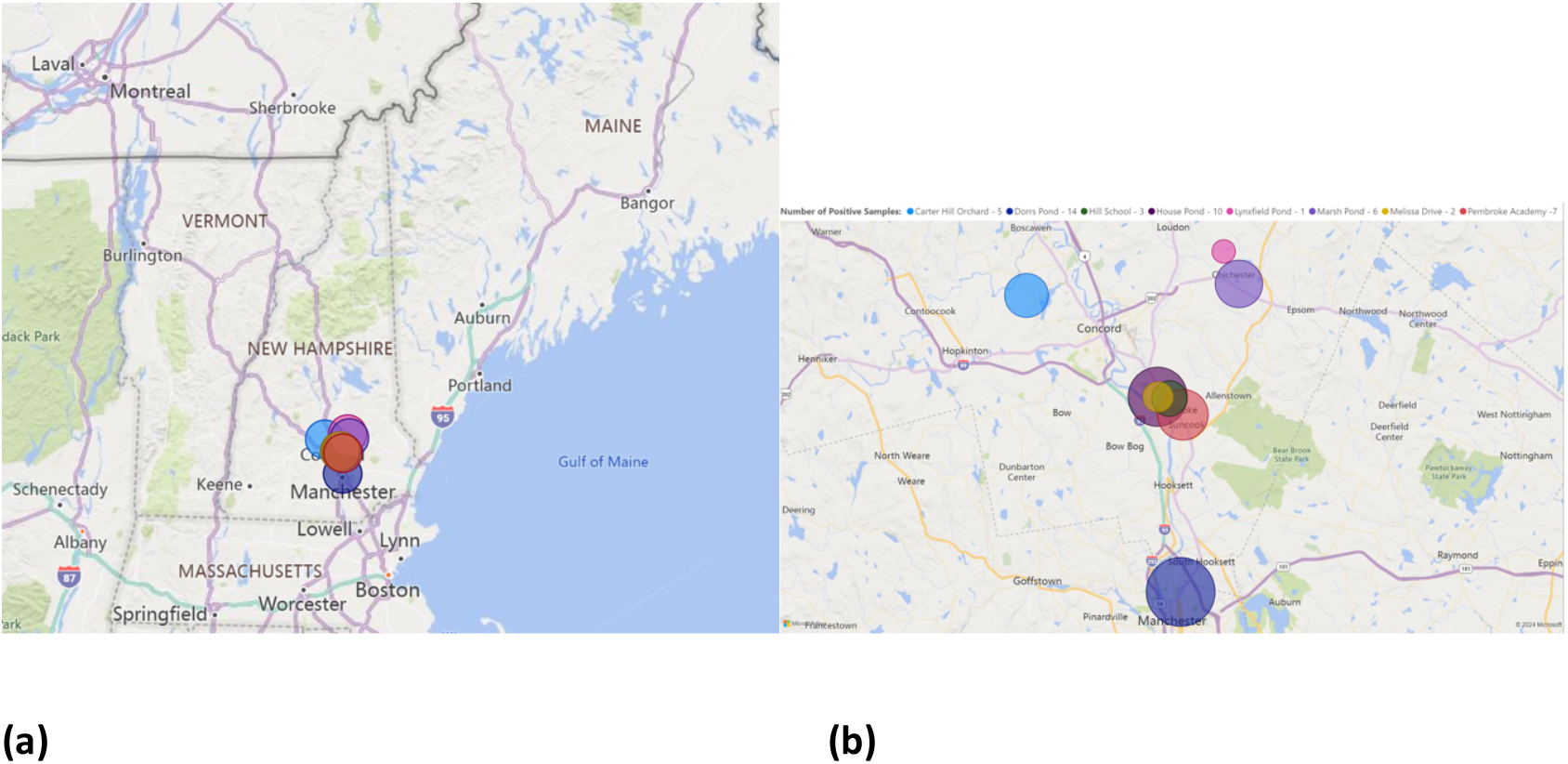
Part (a) shows the location of collecting sites in central and southern New Hampshire, USA as colored circles. Part (b) shows a close-up of the locations of the collecting sites and the color indicates the number of frogs that tested positive for *Bd* at each site.

**Figure 2:**
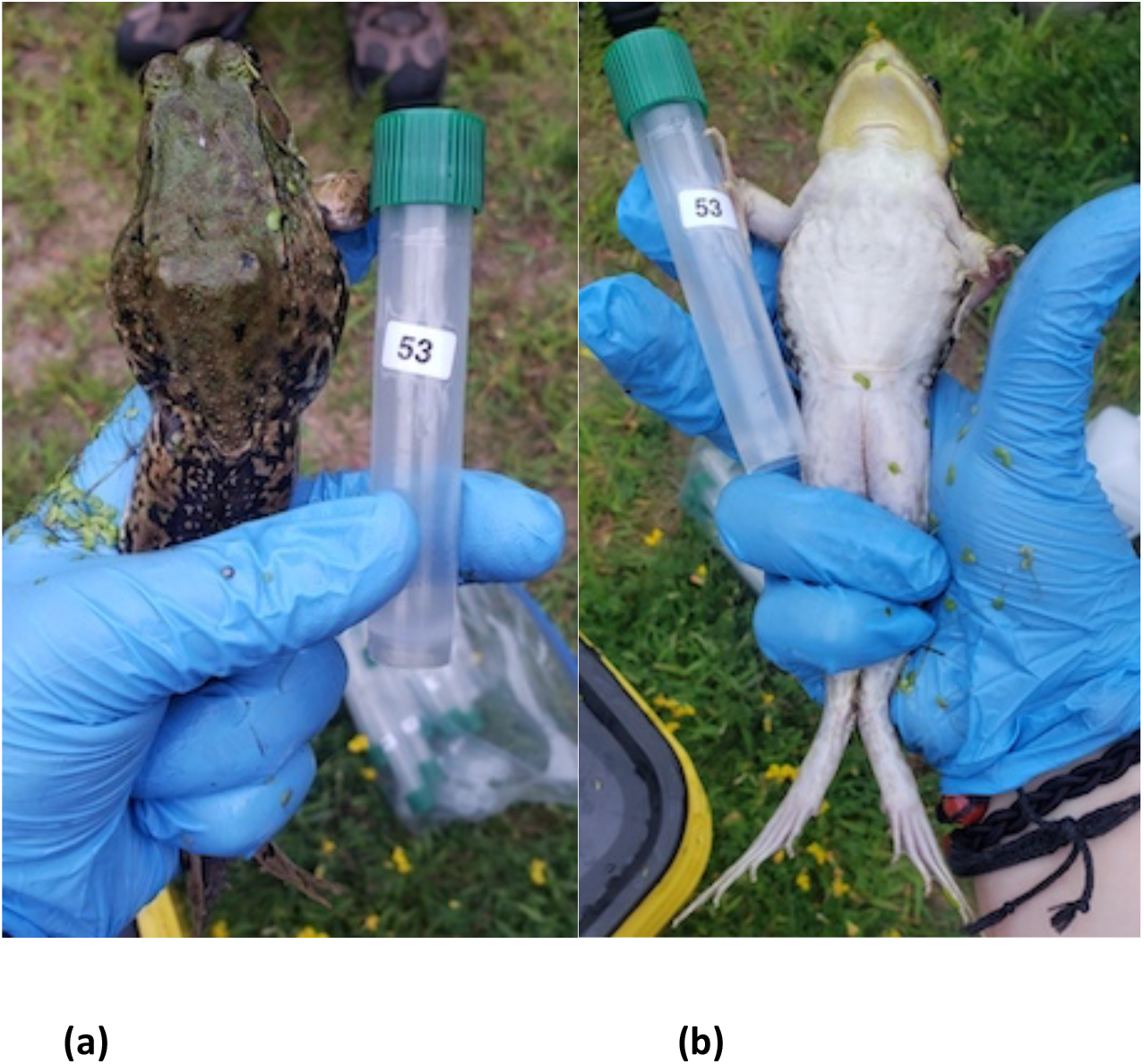
Each collected frog is photographed from both the dorsal (a) and ventral side (b) with its collection vial, for reference. This allows for a digital record and provides another dataset for potential image analysis of the sampled frogs.

After this data was recorded, the tissue swabbing procedure utilized sterile cotton swabs to collect skin cells from the ventral side of the amphibian. This protocol consisted of swabbing the amphibian’s ventral side ten times on either side of the stomach, five times in a horseshoe pattern on the drink patch, ten times on each inner thigh, and twice between each webbing of their hind feet. A single swipe consisted of the swab traveling up and down the length of the area once while lightly twirling the swab. This methodology is used with the goal to collect sloughing skin cells, which is not invasive or harmful to the amphibians. Collection swabs are then placed in a individually labeled plastic 10 mL collection tube with 5-8mL of 70% isopropyl alcohol for DNA preservation. Once the data and skin cell sample were collected, frogs were released into the waterbody they were originally collected from. Filled collection tubes were stored on ice during transport and then stored in a -20°C laboratory freezer until they underwent DNA analysis.

To eliminate the possibility of recapturing and recollecting from the same specimen, each sample location was collected from only once in the 2019 season. Amphibian capture efforts would cease prior to data and tissue collection to further reduce the possibility of duplicate specimen sampling. All reusable field equipment (buckets, nets, boots, etc.) were sterilized in a 5-10% bleach solution and air dried between each sampling location to prevent cross contamination or the possibility of spreading organisms between multiple locations.

### Laboratory Protocols for *Bd* Monitoring

To extract the DNA from the field collected samples, Qiagen DNeasy Blood & Tissue Kit (Qiagen) extraction kits were used. Sample DNA was amplified using GoTaq Hot Start Green MM (Promega) and published primers (Boyle *et al*., 2004). All field-collected samples were tested with positive and negative controls. Positive controls were obtained from the Collection of Zoosporic Eufungi at the University of Michigan. To amplify DNA, samples were placed in a PCR Thermocycler (Biometra TAdvanced) and amplified using a two-minute initial denaturation at 94°C, followed by 40 cycles of a 15 second denaturation step at 94°C, a 30 second annealing step at 60°C, and a one minute, 15 second elongation step at 72°C, preceded by a final extension of five minutes at 72°C. To detect for the presence or absence of *Bd*, we performed agarose gel electrophoresis and visualized samples using an LED light box (Carolina Biological Supply). This is a relatively low-cost method that can be done without a real-time PCR machine and could be performed at many smaller institutions.

### DNA Sequencing

Three samples that tested positive for *Bd* were selected for DNA sequencing. These samples were collected in Pembroke, NH, from two wood frogs (samples 43 and 48) and one bullfrog (sample 172). Samples were chosen based on having a strong band on an agarose gel. These samples were amplified using the Bda1 and Bda2 primers (Annis *et al*., 2004), and then sent for purification and Sanger sequencing to Genewiz (Azenta Life Sciences). The specified primers amplify the ITS1 and ITS2 sections of *Bd* (Annis *et al*., 2004). Once sequences were obtained, we used BLAST to compare our sequences against other sequences deposited on GenBank. Sequences were deposited on GenBank following analysis and assigned accession numbers (Appendix 2).

## Results

### *Bd* Monitoring Effort

In the 2019 monitoring effort, frogs were sampled at eight collection sites across central and southern New Hampshire (Table 1, Fig. 1). Five different species were examined, including bullfrogs, green frogs, spring peepers, wood frogs, and pickerel frogs. We tested a total of 74 frogs, of which 48 tested positive for *Bd* (Table 1). Infections were detected at all collection sites and among all species tested, with the exception of spring peepers (Table 1, Fig. 1). However, it should be noted that only two spring peepers were sampled in 2019.

### DNA Sequencing

Of the three samples tested genetically, at least two different strains were present based on BLAST results. The genetic differences observed are in the ITS (Internal Transcribed Spacers) region, a region that allows for a “fingerprint” to identify different genetic strains and subspecies (Voigt *et al*., 2021; Walker *et al*., 2022). Sample 178 exhibited one deletion and six substitutions, which was the most notable variation from previously documented strains on GenBank (Fig. 3). Figures 4 and 5 show samples 48 and 43 respectively, and each of these are genetically similar to strains previously documented in the literature. Figure 6 shows a BLAST of sample 43 compared to sample 48. Notably, there is a 95% match but still a 5% difference between the two. This indicates that there are likely multiple strains of *Bd* in our New Hampshire populations.

**Figure 3:**
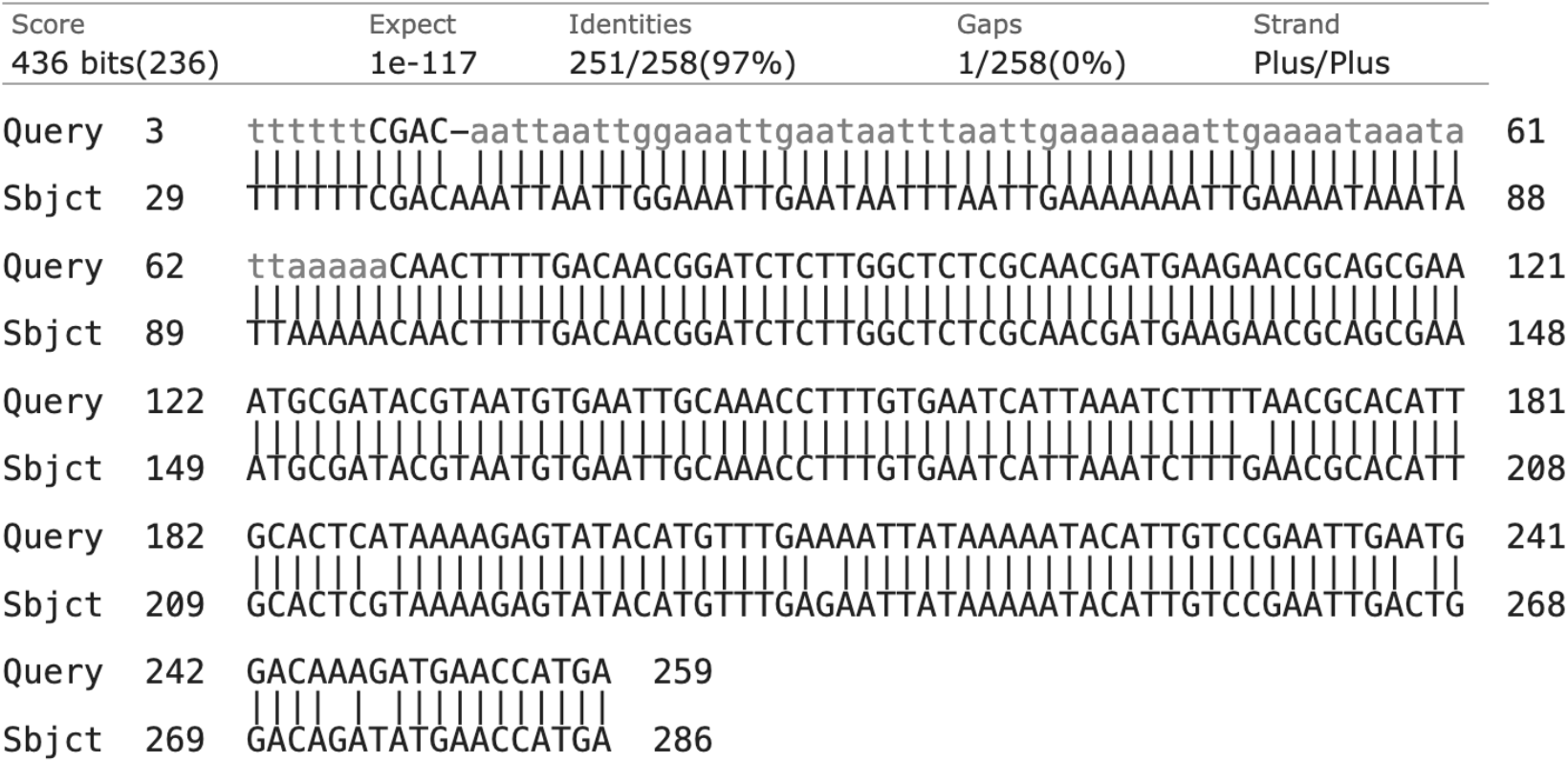
DNA sequence for sample 172 BLAST against MH745069.1 sequence (Vörös *et al*., 2018) with a 97% match. There is a deletion in row one, a substitution in row three and multiple substitutions in rows four and five. Substitutions and deletions can be seen by the absence of vertical line connections. The vertical line connections are only visible on matching, identical base pairs.

**Figure 4:**
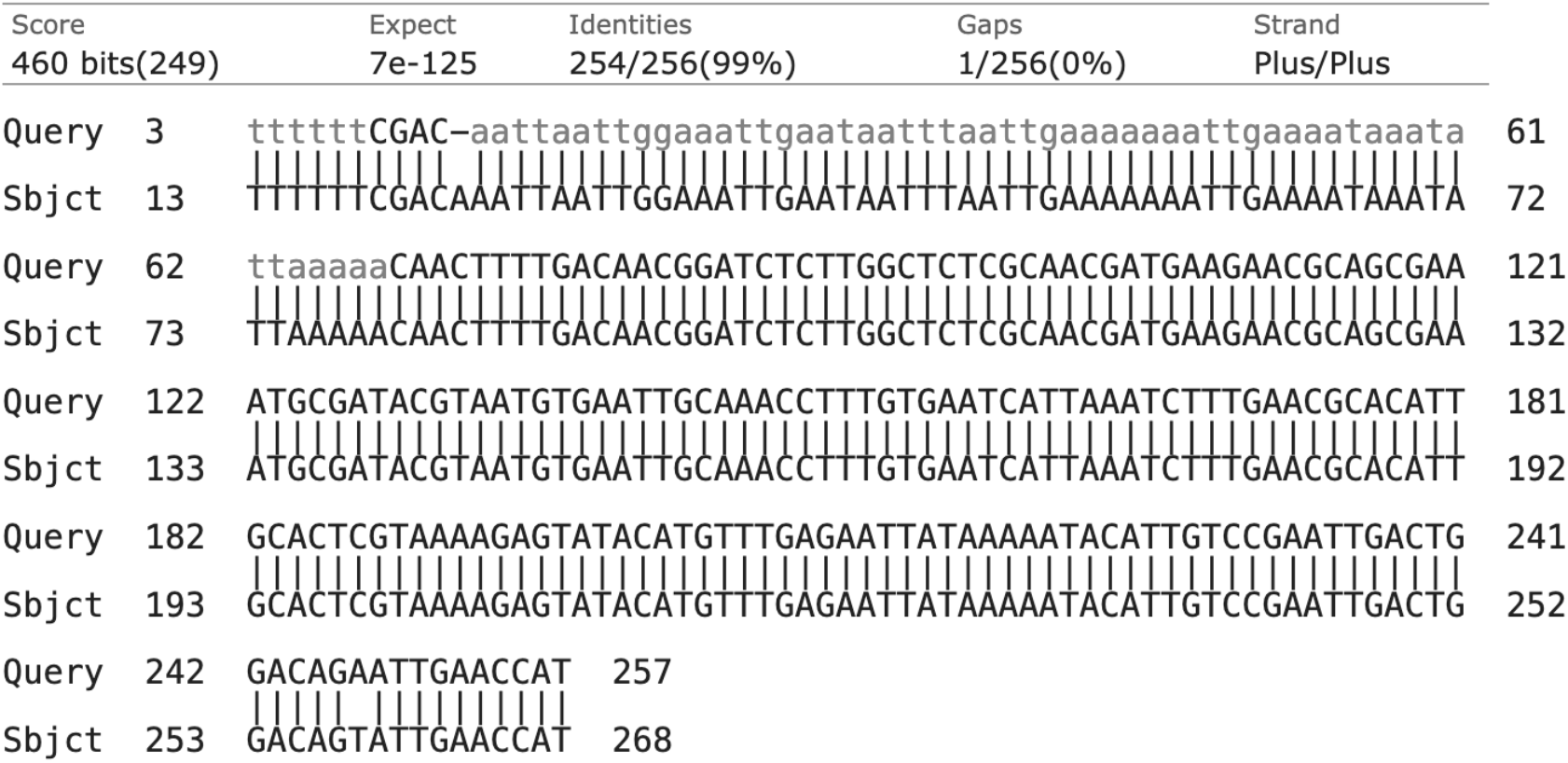
DNA sequence for sample 48 BLAST against FJ229469.1 sequence (Rodriguez *et al*., 2009) with a 99% match. This sequence has one deletion (first row) and one substitution (fifth row).

**Figure 5:**
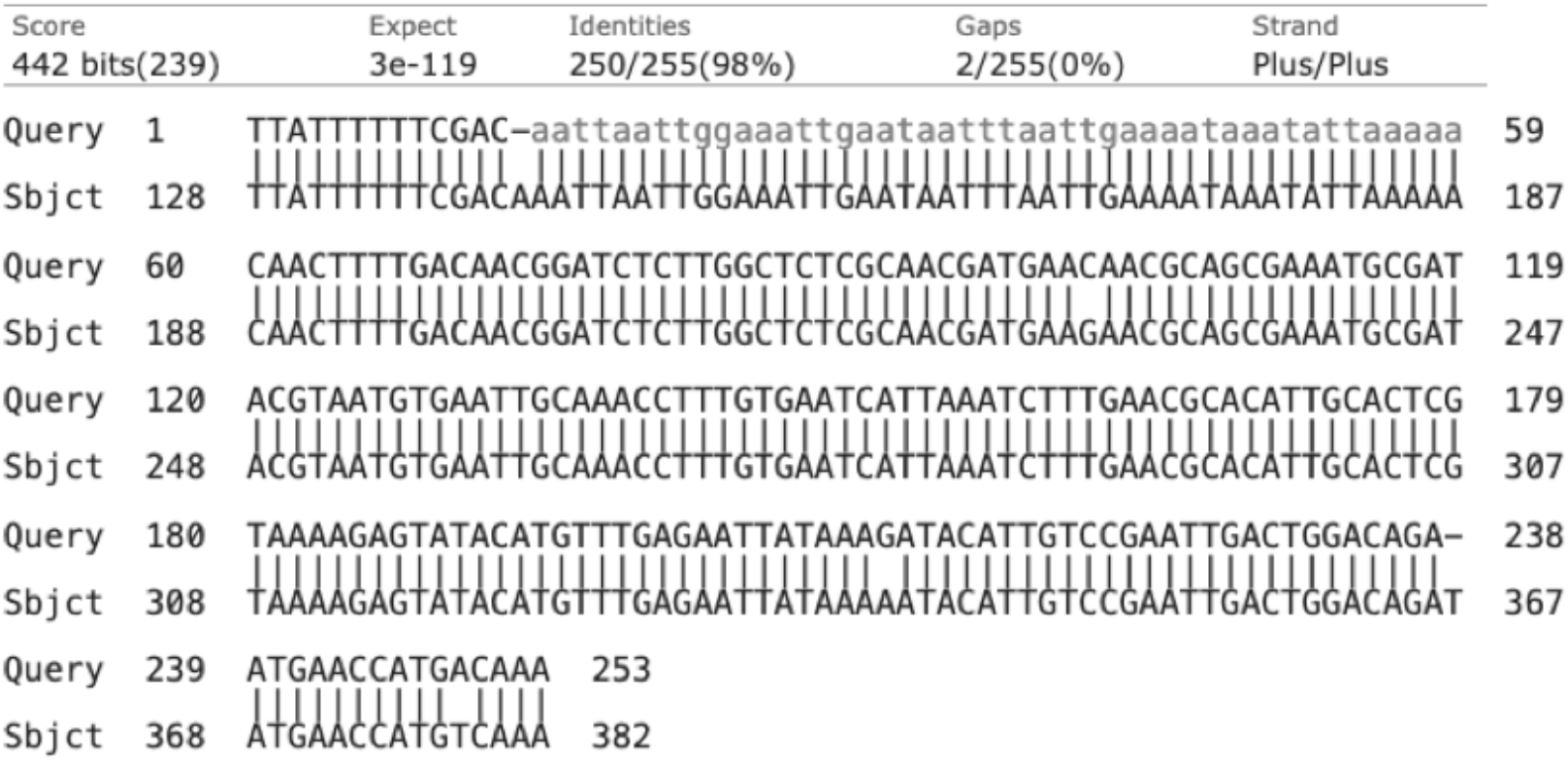
DNA Sequence for sample 43 (lower row) BLAST against JQ582906.1 sequence (upper row) from Schloegel *et al*., (2012). There is a 98% match between the two sequences. Most notably there is one deletions for sample 43 (first row), as well as three substitutions (second, fourth, and fifth row).

**Figure 6:**
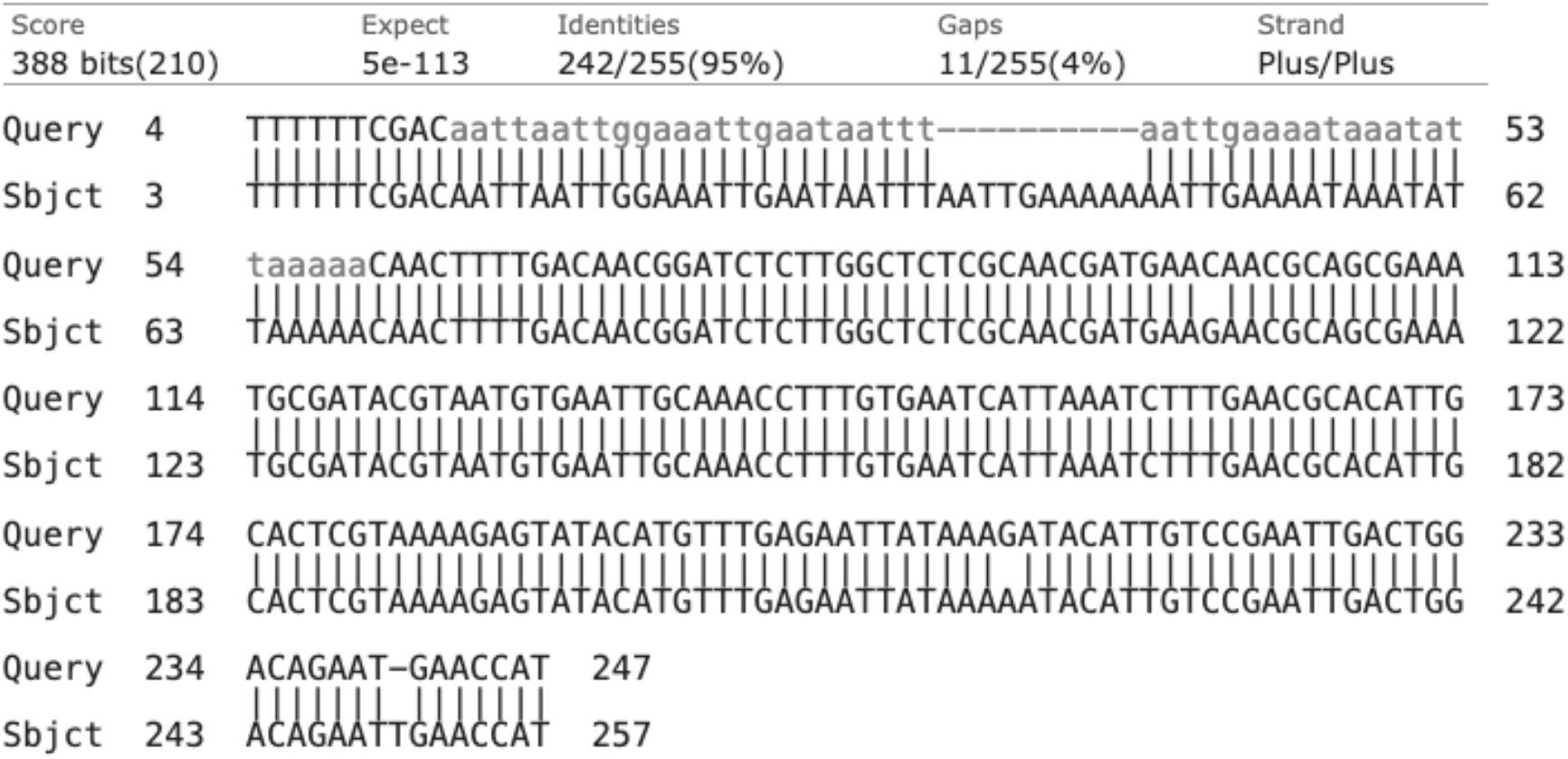
DNA sequence from sample 43 BLAST against DNA sequence from sample 48. The two strains found within our populations are compared to each other to highlight the differences and show that they are distinct strains.

## Discussion

The results of the 2019 efforts of the *Bd* monitoring project showed that nearly 65% of the 74 samples collected tested positive for *Bd*. With such a large percentage of specimens testing positive for chytrid fungus and there being no observations of population decline, this suggests that either the strains in New Hampshire (NH) are less virulent, the frogs have evolved resistance to these strains, or there are environmental factors that are protecting the frogs in NH from succumbing to chytrid fungus. The genetic study results show that in NH, specifically in the town of Pembroke, there are different strains from strains recorded in other areas. These strains were identified by sequencing the ITS, or Internal Transcribed Spacers, of three samples that tested positive for *Bd*. These spacers are unused sections of DNA but they are key to identifying different strains of bacteria, fungi and more, as well as assigning sub-species level of identification (Voigt *et al*., 2021; Walker *et al*., 2022). However, since these are spacer regions, these results alone do not indicate whether the strains in New Hampshire are more or less virulent when compared to other strains. They do indicate previously unidentified strains and further testing can be done to see if there is a link to virulence from these new strains.

Chytrid normally reproduces asexually, but there are documented cases of sexual reproduction in *Bd* (Schloegel *et al*., 2012; Bryne *et al*., 2016); therefore, further examining the genetics of different strains could help to better understand the origin of these new strains and their potential virulence. Genetic variation with sexual reproduction has two mechanisms that can lead to variation, independent assortment of gametes and genetic recombination in the form of crossing over. Further research should be done to better understand the extent of sexual reproduction in *Bd*, and its potential to create new genetic strains.

It is also important to note that while sample 172 appears to be the most different from previously recorded strains, samples 48 and 43 are nearly identical strains to others found on GenBank. This means there is evidence of variance in the area, but also evidence of a single strain mostly unchanged from those found in other regions. It would be worth studying this strain to see what makes it persist in local populations and to better understand how this strain is affecting the health of amphibians in the northeastern United States.

Although there is an increasing amount of work on the genetics of *Bd*, we still know relatively little about how genetics affect virulence (but see Refsnider *et al*., 2015; Bryne *et al*., 2016). Future studies could examine other genes to determine which strains are more virulent or cause greater mortality in various species of frogs. If more areas in the country tested these spacer regions and identified different strains, this could help to identify which strains are less pathogenic to frogs and which are causing the frogs to have increased mortality rates. Additionally, a future study could examine how the changing climate could drive evolution of variation in strains, as well as comparing strains from different climates, such as tropical versus temperate.

Furthermore, this study gives a template for using undergraduate research to provide an ongoing monitoring effort of *Bd* to areas that are understudied. We detected cases of *Bd* at several sites across central and southern New Hampshire. Without our undergraduate research and community-based monitoring effort, these cases would have likely gone undetected. We recommend that more small universities implement *Bd* monitoring programs to better understand the spread and strain diversity of this fungal disease. The methods reported here could even be implemented in many high schools or community science centers and programs. In addition to monitoring *Bd* spread, the monitoring program provides essential training to students in field and molecular techniques. Monitoring for the spread and genetic diversity of this fungus will not only lead to a better understanding of its presence, but it will also help to monitor amphibian populations. Amphibians play a vital role in the health and stability of ecosystems and these monitoring efforts could help us to better protect them. We hope other institutions adopt and implement these techniques to help further understand the spread of *Bd* and provide science training to students and community members.

## Acknowledgements

The authors thank all landowners and business owners who allowed collections on their property. We thank all student workers and community members who dedicated their time to this project, both in the field and in the lab. Special acknowledgements go to Marianne DiTaranto and Julia Hromis for their efforts in the 2019 field collections.

## Author contributions

**Katharine York, Mariah Mitchell**, and **M. Catherine Duryea** conceived of the project and designed the methodology. **Katharine York** and **Mariah Mitchell** led in the field collections. **Tyler Cardoso** and **M. Catherine Duryea** designed the methodology and collected and analyzed data for the DNA sequencing project. **Tyler Cardoso, Ashley Chapdelaine, Yuhui Song**, and **M. Catherine Duryea** collected the laboratory data. **Tyler Cardoso** and **M. Catherine Duryea** led in writing the manuscript. All authors contributed feedback and edits to the manuscript.

## Ethics Statement

The authors confirm that the research detailed here conforms to the ethics of their research institution.

## Conflict of interest

The authors declare no conflicts of interest.

## Notes

### Competing Interest Statement

The authors have declared no competing interest.

