## Supplemental Table 1 for "Undergraduate research project uncovers a novel strain of amphibian chytrid fungus in the Northeastern United States"

**Appendix 1:** Field collection data and PCR results of *Bd* testing. Bolded samples were sent for sequencing.

| Frog # | Sex | Age | Species | Date | Collection Site | Town | Approx. GPS Coord. | Notes | Water Temp (C) | Time | Location Notes | PCR Result |
|---|---|---|---|---|---|---|---|---|---|---|---|---|
| 74 | F | Adult | Bullfrog | 7/23/19 | House Pond | Pem-broke | 43.158661, -71.477785 | Healthy & Active | N/A | 21:05 | Manmade, residential pond | positive |
| 62 | F | Adult | Bullfrog | 7/23/19 | House Pond | Pem-broke | 43.158661, -71.477785 | Healthy | N/A | 21:10 | Manmade, residential pond | positive |
| 98 | F | Adult | Bullfrog | 7/23/19 | House Pond | Pem-broke | 43.158661, -71.477785 | Healthy & Active | N/A | 21:30 | Manmade, residential pond | positive |
| 73 | M | Adult | Bullfrog | 7/23/19 | House Pond | Pem-broke | 43.158661, -71.477785 | Healthy & Active | N/A | 21:45 | Manmade, residential pond | positive, faint band |

| | | | | | | | | | | | | |
|---|---|---|---|---|---|---|---|---|---|---|---|---|
| 77 | M | Adult | Green frog | 7/23/19 | House Pond | Pem-broke | 43.158661, -71.477785 | Healthy | N/A | 21:00 | Manmade, residential pond | positive |
| 79 | M | Adult | Green frog | 7/23/19 | House Pond | Pem-broke | 43.158661, -71.477785 | Healthy & Active | N/A | 21:15 | Manmade, residential pond | positive |
| 91 | M | Adult | Green frog | 7/23/19 | House Pond | Pem-broke | 43.158661, -71.477785 | Healthy | N/A | 21:20 | Manmade, residential pond | negative |
| 76 | M | Adult | Green frog | 7/23/19 | House Pond | Pem-broke | 43.158661, -71.477785 | Active, pinkish marks on groin and feet | N/A | 21:25 | Manmade, residential pond | positive |
| 100 | M | Adult | Green frog | 7/23/19 | House Pond | Pem-broke | 43.158661, -71.477785 | Active, blister on toe, pink | N/A | 21:35 | Manmade, residential pond | negative |

| | | | | | | | | | | | | |
|---|---|---|---|---|---|---|---|---|---|---|---|---|
| | | | | | | | | marks on groin and feet | | | | |
| 70 | M | Adult | Green frog | 7/23/19 | House Pond | Pem-broke | 43.158661, -71.477785 | Healthy | N/A | 21:40 | Manmade, residential pond | positive, faint band |
| 81 | M | Adult | Green frog | 7/23/19 | House Pond | Pem-broke | 43.158661, -71.477785 | Healthy | N/A | 21:50 | Manmade, residential pond | positive |
| 93 | M | Adult | Green frog | 7/23/19 | House Pond | Pem-broke | 43.158661, -71.477785 | Healthy | N/A | 21:55 | Manmade, residential pond | positive |
| 26 | M | Adult | Green frog | 8/17/19 | Dorrs Drainage | Man-chester | 43.012986, -71.454348 | Healthy | 23.889 | 18:30 | Manmade drainage pond | positive |
| 56 | F | Adult | Green frog | 8/17/19 | Dorrs Drainage | Man-chester | 43.012986, -71.454348 | Healthy | 23.889 | 18:30 | Manmade drainage pond | positive, faint band |

| | | | | | | | | | | | | |
|---|---|---|---|---|---|---|---|---|---|---|---|---|
| 53 | F | Adult | Green frog | 8/17/19 | Dorrs Drainage | Man-chester | 43.012986, -71.454348 | Healthy | 23.889 | 18:30 | Manmade drainage pond | positive |
| 66 | F | Adult | Bullfrog | 8/17/19 | Dorrs Pond - East Side | Man-chester | 43.014053, -71.453705 | Healthy | 23.333 | 19:40 | Natural, managed pond | positive, faint band |
| 64 | F | Adult | Green frog | 8/17/19 | Dorrs Pond - East Side | Man-chester | 43.014053, -71.453705 | Healthy | 23.333 | 19:53 | Natural, managed pond | positive |
| 88 | F? | Juv | Green frog | 8/17/19 | Dorrs Pond - East Side | Man-chester | 43.014053, -71.453705 | Healthy | 23.333 | 19:58 | Natural, managed pond | positive |
| 161 | F | Adult | Green frog | 8/17/19 | Dorrs Pond - East Side | Man-chester | 43.014053, -71.453705 | Healthy | 23.333 | 20:05 | Natural, managed pond | positive |
| 72 | F | Adult | Green frog | 8/17/19 | Dorrs Pond - East Side | Man-chester | 43.014053, -71.453705 | Healthy | 23.333 | 20:07 | Natural, managed pond | negative |

| | | | | | | | | | | | | |
|---|---|---|---|---|---|---|---|---|---|---|---|---|
| 160 | F | Y. Adult | Green frog | 8/17/19 | Dorrs Pond - East Side | Man-chester | 43.014053, -71.453705 | Healthy | 23.333 | 20:12 | Natural, managed pond | positive |
| 78 | M | Adult | Green frog | 8/17/19 | Dorrs Pond - East Side | Man-chester | 43.014053, -71.453705 | Healthy | 23.333 | 20:15 | Natural, managed pond | negative |
| 158 | F | Adult | Green frog | 8/17/19 | Dorrs Pond - East Side | Man-chester | 43.014053, -71.453705 | Healthy | 23.333 | 20:19 | Natural, managed pond | negative |
| 89 | F | Juv | Green frog | 8/17/19 | Dorrs Pond - East Side | Man-chester | 43.014053, -71.453705 | Healthy | 23.333 | 20:25 | Natural, managed pond | negative |
| 22 | F | Y. Adult | Green frog | 8/25/19 | LynxField Pond | Chi-chester | 43.266915, -71.410354 | Healthy | 23 | 16:30 | Natural water body | Negative |
| 46 | F | Y. Adult | Green frog | 8/25/19 | LynxField Pond | Chi-chester | 43.266915, -71.410354 | Healthy | 23 | 16:30 | Natural water body | Negative |

| | | | | | | | | | | | | |
|---|---|---|---|---|---|---|---|---|---|---|---|---|
| 14 | F | Adult | Green frog | 8/25/19 | LynxField Pond | Chi- chester | 43.266915, - 71.410354 | Healthy, thighs reddish | 23 | 16:30 | Natural water body | positive |
| 42 | M | Adult | Green frog | 8/25/19 | Marsh Pond | Chi- chester | 43.242743, - 71.394715 | Healthy | 25 | 13:30 | Manmade and Natural waterbody (combo) | positive |
| 9 | F | Adult | Green frog | 8/25/19 | Marsh Pond | Chi- chester | 43.242743, - 71.394715 | Pink patch under right foot | 25 | 13:30 | Manmade and Natural waterbody (combo) | positive |
| 8 | M | Adult | Green frog | 8/25/19 | Marsh Pond | Chi- chester | 43.242743, - 71.394715 | Blood on thighs, red feet | 25 | 13:30 | Manmade and Natural waterbody (combo) | positive |
| 10 | ? | Y. Adult | Pickerel Frog | 8/25/19 | Marsh Pond | Chi- chester | 43.242743, - 71.394715 | left foot missing | 25 | 13:30 | Manmade and Natural | positive |

| | | | | | | | | | | | | |
|---|---|---|---|---|---|---|---|---|---|---|---|---|
| | | | | | | | | | | | waterbody (combo) | |
| 17 | F | Y. Adult | Pickerel Frog | 8/25/19 | Marsh Pond | Chi-chester | 43.242743, -71.394715 | Mark on thighs | 25 | 13:30 | manmade and Natural waterbody (combo) | positive |
| 37 | F | Y. Adult | Pickerel Frog | 8/25/19 | Marsh Pond | Chi-chester | 43.242743, -71.394715 | N/A | 25 | 13:30 | manmade and Natural waterbody (combo) | negative |
| 25 | ? | Y. Adult | Pickerel Frog | 8/25/19 | Marsh Pond | Chi-chester | 43.242743, -71.394715 | Dark Spots on Chest | 25 | 13:30 | manmade and Natural waterbody (combo) | positive |
| 7 | F | Adult | Green frog | 9/1/19 | Hill School | Pem-broke | 43.157363, -71.465946 | Healthy | 17.222 | 19:30 | N/A | negative |
| 18 | F | Adult | Green frog | 9/1/19 | Hill School | Pem-broke | 43.157363, -71.465946 | Healthy | 17.222 | 19:30 | N/A | positive |

| | | | | | | | | | | | | |
|---|---|---|---|---|---|---|---|---|---|---|---|---|
| 16 | F | Adult | Green frog | 9/1/19 | Hill School | Pem-broke | 43.157363, -71.465946 | Healthy | 17.222 | 19:30 | N/A | positive |
| 31 | F | Adult | Green frog | 9/1/19 | Hill School | Pem-broke | 43.157363, -71.465946 | Healthy | 17.222 | 19:30 | N/A | positive |
| 5 | F | Y. Adult | Green frog | 9/1/19 | Hill School | Pem-broke | 43.157363, -71.465946 | Healthy | 17.222 | 19:30 | N/A | negative |
| 11 | F | Adult | Bullfrog | 9/2/19 | PA Pond | Pem-broke | 43.145273, -71.452083 | Healthy | 15.556 | 17:45 | Note - rainy day | positive, faint band |
| 35 | F | Adult | Bullfrog | 9/2/19 | PA Pond | Pem-broke | 43.145273, -71.452083 | Healthy | 15.556 | 18:00 | Note - rainy day | positive, faint band |
| 2 | F | Adult | Bullfrog | 9/2/19 | PA Pond | Pem-broke | 43.145273, -71.452083 | Losing a lot of skin | 15.556 | 18:00 | Note - rainy day | negative |
| 36 | F | Y. Adult | Green frog | 9/2/19 | PA Pond | Pem-broke | 43.145273, -71.452083 | Healthy | 15.556 | 18:00 | Note - rainy day | Negative |
| 164 | F | Adult | Bullfrog | 9/6/19 | PA Pond | Pem-broke | 43.145273, -71.452083 | None | 20 | 18:00 | Nighttime/after dark frogging | positive, faint band |

| | | | | | | | | | | | | |
|---|---|---|---|---|---|---|---|---|---|---|---|---|
| 165 | F | Adult | Bullfrog | 9/6/19 | PA Pond | Pem-broke | 43.145273, -71.452083 | None | 20 | 18:00 | nighttime/after dark frogging | negative |
| 171 | F | Y. Adult | Bullfrog | 9/6/19 | PA Pond | Pem-broke | 43.145273, -71.452083 | skinny | 20 | 18:00 | nighttime/after dark frogging | positive, faint band |
| 175 | F | Adult | Bullfrog | 9/6/19 | PA Pond | Pem-broke | 43.145273, -71.452083 | None | 20 | 18:00 | nighttime/after dark frogging | Negative |
| 172 | F | Adult | Bullfrog | 9/6/19 | PA Pond | Pem-broke | 43.145273, -71.452083 | skinny | 20 | 18:00 | nighttime/after dark frogging | positive |
| 1 | F | Adult | Bullfrog | 9/6/19 | PA Pond | Pem-broke | 43.145273, -71.452083 | None | 20 | 18:00 | nighttime/after dark frogging | positive, faint band |
| 168 | F | Adult | Green frog | 9/6/19 | PA Pond | Pem-broke | 43.145273, -71.452083 | None | 20 | 18:00 | nighttime/after dark frogging | Negative |

| | | | | | | | | | | | | |
|---|---|---|---|---|---|---|---|---|---|---|---|---|
| 155 | F | Adult | Green frog | 9/6/19 | PA Pond | Pem-broke | 43.145273, -71.452083 | None | 20 | 18:00 | nighttime/after dark frogging | Negative |
| 39 | F | Adult | Green frog | 9/6/19 | PA Pond | Pem-broke | 43.145273, -71.452083 | None | 20 | 18:00 | nighttime/after dark frogging | Negative |
| 174 | F | Adult | Green frog | 9/6/19 | PA Pond | Pem-broke | 43.145273, -71.452083 | skinny | 20 | 18:00 | nighttime/after dark frogging | positive, faint band |
| 141 | F | Adult | Bullfrog | 9/29/19 | Carter Hill Orchard | Con-cord | 43.234157, -71.612004 | Healthy | 19.778 | 18:19 | Manmade irrigation pond | positive, faint band |
| 131 | F | Adult | Bullfrog | 9/29/19 | Carter Hill Orchard | Con-cord | 43.234157, -71.612004 | None | 19.778 | 18:20 | Manmade irrigation pond | positive |
| 128 | F | Adult | Bullfrog | 9/29/19 | Carter Hill Orchard | Con-cord | 43.234157, -71.612004 | Blister on feet, took | 19.778 | 18:23 | Manmade irrigation pond | positive |

| | | | | | | | | | | | | |
|---|---|---|---|---|---|---|---|---|---|---|---|---|
| | | | | | | | | excess skin for testing | | | | |
| 132 | F | Adult | Bullfrog | 9/29/19 | Carter Hill Orchard | Con-cord | 43.234157, -71.612004 | None | 19.778 | 18:24 | Manmade irrigation pond | positive, faint band |
| 152 | F | Adult | Green Frog | 9/29/19 | Carter Hill Orchard | Con-cord | 43.234157, -71.612004 | Red marks on bottom of feet | 19.778 | 18:14 | Manmade irrigation pond | negative |
| 169 | F | Adult | Green Frog | 9/29/19 | Carter Hill Orchard | Con-cord | 43.234157, -71.612004 | Healthy | 19.778 | 18:24 | Manmade irrigation pond | positive |
| 149 | F | Adult | Bullfrog | 10/6/19 | Dorrs Pond - East Side | Man-chester | 43.014053, -71.453705 | None | 11.889 | 16:40 | Natural, managed pond | positive |

| | | | | | | | | | | | | |
|---|---|---|---|---|---|---|---|---|---|---|---|---|
| 145 | F | Y. Adult | Bullfrog | 10/6/19 | Dorrs Pond - East Side | Man-chester | 43.014053, -71.453705 | None | 11.889 | 16:40 | Natural, managed pond | positive |
| 139 | ? | Juv | Bullfrog | 10/6/19 | Dorrs Pond - East Side | Man-chester | 43.014053, -71.453705 | Lump on back hips | 11.889 | 16:40 | Natural, managed pond | negative |
| 126 | ? | Y. Adult | Green Frog | 10/6/19 | Dorrs Pond - East Side | Man-chester | 43.014053, -71.453705 | Found dead and swollen, looked healthy/no sign of injury | | 16:40 | Natural, managed pond | negative |
| 137 | M | Adult | Green Frog | 10/6/19 | Dorrs Pond - East Side | Man-chester | 43.014053, -71.453705 | None | 11.889 | 16:40 | Natural, managed pond | positive |
| 167 | F | N/A | Green Frog | 10/6/19 | Dorrs Pond - East Side | Man-chester | 43.014053, -71.453705 | None | 11.889 | 16:40 | Natural, managed pond | positive |

| | | | | | | | | | | | | |
|---|---|---|---|---|---|---|---|---|---|---|---|---|
| 138 | F | Y. Adult | Green Frog | 10/6/19 | Dorrs Pond - East Side | Man-chester | 43.014053, -71.453705 | None | 11.889 | 16:40 | Natural, managed pond | positive |
| 153 | F | Y. Adult | Green Frog | 10/6/19 | Dorrs Pond - East Side | Man-chester | 43.014053, -71.453705 | None | 11.889 | 16:40 | Natural, managed pond | positive |
| 45 | F | Adult | Green Frog | 10/7/19 | Melissa Drive | Pem-broke | 43.158510, -71.477469 | Healthy | 17.222 | 21:00 | Rainy Night/ Amphibian Migration | negative |
| 42 | ? | Adult | Spring Peeper | 10/7/19 | Melissa Drive | Pem-broke | 43.158510, -71.477469 | Healthy | 17.222 | 21:00 | Rainy Night/ Amphibian Migration | negative |
| 47 | ? | Adult | Spring Peeper | 10/7/19 | Melissa Drive | Pem-broke | 43.158510, -71.477469 | Healthy | 17.222 | 21:00 | Rainy Night/ Amphibian Migration | negative |
| 43 | ? | Adult | Wood Frog | 10/7/19 | Melissa Drive | Pem-broke | 43.158510, -71.477469 | Healthy | 17.222 | 21:00 | Rainy Night/ Amphibian Migration | positive |

| | | | | | | | | | | | | |
|---|---|---|---|---|---|---|---|---|---|---|---|---|
| 50 | ? | Adult | Wood frog | 10/7/19 | Melissa Drive | Pem-broke | 43.158510, -71.477469 | Healthy | 17.222 | 21:00 | Rainy Night/ Amphibian Migration | negative |
| 49 | ? | Adult | Wood frog | 10/7/19 | Melissa Drive | Pem-broke | 43.158510, -71.477469 | Healthy | 17.222 | 21:00 | Rainy Night/ Amphibian Migration | negative |
| 48 | ? | Adult | Wood frog | 10/7/19 | Melissa Drive | Pem-broke | 43.158510, -71.477469 | Had excess fluid around mouth | 17.222 | 21:00 | Rainy Night/ Amphibian Migration | positive |
