## Supplemental Table 2 for "Undergraduate research project uncovers a novel strain of amphibian chytrid fungus in the Northeastern United States"

**Appendix 2:** GenBank Accession numbers for DNA sequences collected in this study.

| **Sample ID** | **Accession #** |
|---|---|
| Sample 172 | PQ231056 |
| Sample 48 | PQ231057 |
| Sample 43 | PQ231058 |
